# Superovulation does not alter calcium oscillations following fertilization

**DOI:** 10.1101/2021.08.23.457392

**Authors:** Virginia Savy, Paula Stein, Min Shi, Carmen J. Williams

**Affiliations:** Reproductive and Developmental Biology Laboratory, National Institute of Environmental Health Sciences, National Institutes of Health, Research Triangle Park, NC 27709, USA; Biostatistics & Computational Biology Branch, National Institute of Environmental Health Sciences, National Institutes of Health, Research Triangle Park, NC 27709, USA

**Keywords:** Superovulation, oocyte, mouse, calcium oscillations, egg activation

## Abstract

Superovulation is a common approach to maximize the number of eggs available for either clinical assisted reproductive technologies or experimental animal studies. This procedure provides supraphysiological amounts of gonadotropins to promote continued growth and maturation of ovarian follicles that otherwise would undergo atresia. There is evidence in mice, cows, sheep and humans that superovulation has a detrimental impact on the quality of the resulting ovulated eggs or embryos. Here we tested the hypothesis that eggs derived from superovulation have a reduced capacity to support calcium oscillations following fertilization, which is a critical factor in the success of embryo development. Eggs were obtained from mice that were either naturally cycling or underwent a standard superovulation protocol. Naturally cycling mice were mated to vasectomized males and vaginal plugs were checked to assure ovulation had occurred. The superovulated mice were also mated to vasectomized males for consistency of treatment across groups. The eggs were fertilized in vitro while undergoing monitoring of calcium oscillatory patterns. There were no differences in any measures of calcium oscillatory behavior, including length of the first oscillation, area under the curve of calcium signal, or frequency or persistence of oscillations. These findings indicate that superovulation does not disrupt calcium signaling at fertilization, supporting the use of this method for both clinical and experimental purposes.

## Introduction

Fertilization represents the union of two terminally differentiated gametes to form a single embryo capable of developing into a unique individual. Gamete fusion is only the beginning of the process of embryo development, but it sets in motion a series of events collectively termed “egg activation” that change the unified gametes into a totipotent embryo capable of becoming a healthy individual. The key event of egg activation that triggers development in all species is a rise in the cytoplasmic calcium level (Kashir et al., 2013). What is unusual about mammals is that following the initial calcium rise, they go on to have a series of oscillations in cytoplasmic calcium levels that persist for several hours after fertilization. These calcium oscillations drive downstream events of egg activation including exocytosis of cortical granules, activation of calcium/calmodulin-dependent protein kinase II-gamma, resumption of the cell cycle, and pronucleus formation, all of which are essential for initiating proper embryo development (Stein et al., 2020). In the mouse, an inappropriate pattern of calcium oscillations following fertilization is associated with reductions in implantation efficiency, reduced development to term, and abnormalities in offspring growth (Ozil and Huneau, 2001; Ducibella et al., 2002; Ozil et al., 2005, 2006).

The pattern of calcium oscillations at fertilization is modulated by the amount of PLCζ released by the sperm but also by many factors intrinsic to the fertilized egg (Stein et al., 2020). Calcium oscillations depend on the amount of calcium in endoplasmic reticulum (ER) stores as well as egg factors that regulate how quickly calcium is released from the ER, cleared from the cytoplasm, and then pumped back into the ER such that sufficient calcium stores are available for the next calcium release event. Calcium homeostasis is regulated by the activities of calcium pumps and ion channels (Berridge et al., 2003). Sarco-endoplasmic reticulum calcium-ATPases pump calcium back into the ER, and plasma membrane calcium-ATPase pumps clear calcium from the cytoplasm by extruding it across the plasma membrane. These pumps depend on mitochondrial production of ATP. Calcium influx channels support calcium entry into the cytoplasm down a concentration gradient from the extracellular milieu; calcium influx is necessary for complete refilling of ER stores and continuation of calcium oscillations (Igusa and Miyazaki, 1983; Kline and Kline, 1992). Together, these activities allow the egg to tolerate the release of large amounts of calcium from the ER and to refill ER stores once the calcium release event has been completed.

All the molecular components needed to support calcium homeostasis, egg activation and preimplantation embryo development, such as stored proteins, nucleic acids, and energy substrates, are generated during the oocyte growth phase. In a normal estrous cycle, ovarian follicles are recruited into the growing pool independent of gonadotropin hormones until the secondary follicle stage, when the oocyte is surrounded by several layers of granulosa cells (Strauss III and Williams, 2017). Follicle stimulating hormone (FSH), secreted from the pituitary, initiates further follicle development to the antral stage, when fluid begins to collect between the granulosa cells. FSH and luteinizing hormone (LH) together promote continued survival and development of antral follicles toward the preovulatory stage (Sullivan et al., 1999)(Sullivan et al., 1999)(Chun et al., 1996; Sullivan et al., 1999). When FSH levels decline in response to estradiol-mediated negative feedback on the pituitary, antral follicles instead will undergo atresia. A select few “dominant” follicles continue to grow despite declining FSH levels. These follicles are either more sensitive to the available FSH or have developed sufficient responsiveness to LH to continue their developmental trajectory (Sullivan et al., 1999). As a result, a species-specific number of preovulatory follicles forms during each estrous cycle.

Successful superovulation was first reported almost 100 years ago, when whole pituitary tissue from donor animals was injected intramuscularly into rats and mice and resulted in either ovaries containing very large numbers of normal-appearing follicles or oviducts containing large numbers of ovulated eggs (Smith and Engle, 1927). Although the exact procedure has been modified extensively since that time, superovulation is commonly utilized to maximize the number of ovulated eggs available for either experimental purposes or for clinical use within assisted reproduction cycles. Superovulation works by providing high amounts of exogenous gonadotropin hormone so that non-dominant late secondary and early antral follicles, which are sensitive to gonadotropin stimulation and would otherwise be destined to undergo atresia, are instead maintained in the growing follicle pool and form preovulatory follicles. The preovulatory follicles can be allowed to ovulate spontaneously or induced to ovulate by administration of chorionic gonadotropin, which binds to the LH receptor and begins the ovulatory signaling cascade.

There is evidence in mice, cows, sheep and humans that superovulation has a detrimental impact on the quality of the resulting ovulated eggs or embryos (Moor et al., 1985; Hyttel et al., 1986; Yun et al., 1989; Assey et al., 1994; Blondin et al., 1996; Van der Auwera, 2001; Baart et al., 2007; Fortier et al., 2008; Market-Velker et al., 2010; Lee et al., 2017). These findings could be explained by many factors, including accelerated oocyte growth, rescue of abnormal oocytes from atresia, and recruitment for ovulation of non-mature oocytes, all of which could lead to eggs that lack the maternally derived stores needed for developmental competence. Because calcium oscillatory patterns depend heavily on maternal components that regulate calcium homeostasis, we hypothesized that superovulation results in eggs that have abnormal patterns of calcium oscillations following fertilization. Instead, we found no differences in any parameters of calcium oscillatory behavior following in vitro fertilization when comparing spontaneously ovulated and superovulated eggs.

## Methods

### Animals and superovulation

C57BL/6J females (3-6 weeks old), C57BL/6J vasectomized males (2-6 months old), and B6SJLF1/J males (4-6 months old) were obtained from The Jackson Laboratory (Bar Harbor, ME).

Metaphase II-arrested (MII) eggs were collected from the oviducts of naturally cycling, 6-week-old females following overnight breeding to vasectomized males; vaginal plugs were checked to assure ovulation had occurred. For superovulation, 3-week-old females were primed by intraperitoneal injection of 5 IU of equine chorionic gonadotropin (Lee Biosolutions, Maryland Heights, MO) followed 46–48 h later by 5 IU human chorionic gonadotropin (Sigma Aldrich, St. Louis, MO). The superovulated females were also bred overnight to vasectomized males for consistency across both treatment groups; only vaginal plug-positive females were used.

All mice were sacrificed by CO_2_ asphyxiation and cervical dislocation. All procedures involving mice were conducted in accordance with National Institute of Environmental Health Sciences guidelines under approved animal care and use protocols.

### In vitro fertilization and calcium imaging

Eggs were collected at 9:00 am on the day the vaginal plug was detected, which corresponded to 14 h after hCG injection of superovulated females. Minimal Essential Medium with Hepes (Thermo Fisher, Waltham, MA) containing 0.1% PVA and 0.1% hyaluronidase (Sigma, St. Louis, MO) was used for egg collection and cumulus cell removal. In each biological replicate, eggs from 3-5 mice, either superovulated or naturally cycling, were pooled and treated with acidic Tyrode’s solution (pH 1.6) to remove the zona pellucida. The zona-free eggs were allowed to recover in KSOM medium (Millipore Sigma, Burlington, MA) for 30 min. The eggs were then loaded with the calcium indicator Fura-2 AM (5 μM; Thermo Fisher) for 30 min in KSOM containing 0.02% pluronic F-127 (Thermo Fisher). After being washed in fresh KSOM medium, eggs from superovulated or naturally cycling mice were adhered side by side to Cell-Tak-treated (Thermo Fisher) glass-bottom dishes (MatTek, Ashland, MA) in 150 µl of BSA-free KSOM (Millipore Sigma). Forty-five µl of human tubal fluid medium (HTF; Millipore Sigma) containing 4 mg/ml BSA (HTF-BSA) were added to the drop to achieve a final concentration, once sperm were added, of 1 mg/ml BSA. The drop was covered with mineral oil and the dish placed on the microscope stage in a humidified atmosphere of 5% CO_2_ in room air for intracellular calcium monitoring.

Motile sperm were isolated from the epididymides of adult B6SJLF1/J males in HTF-BSA, via the swim-up method. Briefly, the epididymis and vas deferens were dissected and placed in 500 µl HTF-BSA covered with mineral oil. Under a dissection microscope, the tissue was cut several times and cultured for 15 minutes at 37 °C to allow the motile sperm to swim out. The tissue was removed and 150 µl of swim out sperm were carefully placed at the bottom of a clean tube containing 850 µl of HTF-BSA. The sperm were cultured for 60 min at 37°C to allow the motile sperm to swim up and capacitate. Sperm from the top layer of the culture medium were carefully transferred to a pre-warmed tube and the final sperm concentration was determined using a hemocytometer. Sperm were diluted in HTF-BSA to 10^6^ sperm/ml, then 5 µl were added to the IVF dish (195 µl volume) to achieve a final concentration of 25,000 sperm/ml.

Cytosolic calcium was measured by recording the intensity of fluorescence induced by 340 nm and 380 nm excitation and calculating the F340/F380 ratio. Ratio images were recorded every 7.5 sec using a Hamamatsu ORCA-Flash4.0 LT+ digital camera (Hamamatsu, Bridgewater, NJ) attached to a Nikon Ti inverted microscope with a Nikon S Fluor 20x/0.75 NA objective (Nikon Instruments, Melville, NY) and a Lambda 10-B Optical Filter Changer (Sutter Instruments, Novato, CA). Nikon NIS-Elements software was used for data acquisition and visualization. All experiments were performed at 37°C in a humidified atmosphere of 5% CO_2_, in an Okolab stage microenvironmental chamber enclosed in a microscope cage incubator (Okolab, Ambridge, PA).

A total of 4 biological replicates and 8 technical replicates were analyzed, including 92 eggs from superovulated females and 68 eggs from naturally cycling females. Eggs of each group were fertilized and imaged at the same time, in the same medium drop, for each experimental replicate.

### Data analysis and statistical tests

Custom R functions were developed for automated analysis of calcium imaging data and is available at https://www.niehs.nih.gov/research/atniehs/labs/assets/docs/q_z/savy_rscript.zip. The R-Script was used to determine the time to the first calcium transient, length of the first oscillation, area under the curve (AUC), number of oscillations for 60 and 120 minutes from the starting point of the first transient, number of oscillations per 10 minutes and persistence of oscillations for 120 minutes. Statistical tests were performed using GraphPad Prism. Because the data were not normally distributed, the time to the first transient, length of the first transient, AUC, and oscillation frequency were analyzed using a Mann–Whitney U-test. Data regarding oscillation persistence were analyzed using the Log-rank (Mantel-Cox) test.

## Results

We chose to use the inbred mouse line, C57BL/6, to test whether there were differences in sperm-induced calcium oscillatory patterns between eggs from naturally cycling (NC) and superovulated (SOV) mice. This choice was driven in large part because knockout facilities utilize this strain for the generation of the majority of their animal models. Furthermore, many transgenic and knockout lines are uniquely available on this background strain.

SOV and NC eggs were fertilized in vitro and the sperm-induced calcium oscillatory pattern of eggs in each group were compared. To ensure that the eggs in the two groups were fertilized and cultured under exactly the same conditions, the eggs were adhered side by side in the same medium drop for fertilization and monitoring of calcium oscillatory patterns. There were no obvious differences in the patterns of calcium oscillations (Fig. 1A) or the time until fertilization (Fig. 1B) between SOV and NC eggs. The length of the first transient, which reflects the amount of calcium in endoplasmic reticulum stores, was not different between groups (Fig. 1C). A detailed analysis of additional parameters was performed including the percentage of eggs that continued to display calcium oscillations during 120 min (persistence of oscillations), oscillation frequency, and the area under the curve of calcium signal for the first 60 min following fertilization. No statistically significant differences in any of the parameters analyzed were found between groups (Fig. 1D, 1E, 1F). These findings suggest that superovulation does not impact calcium oscillatory patterns to any measurable degree.

**Figure 1.**
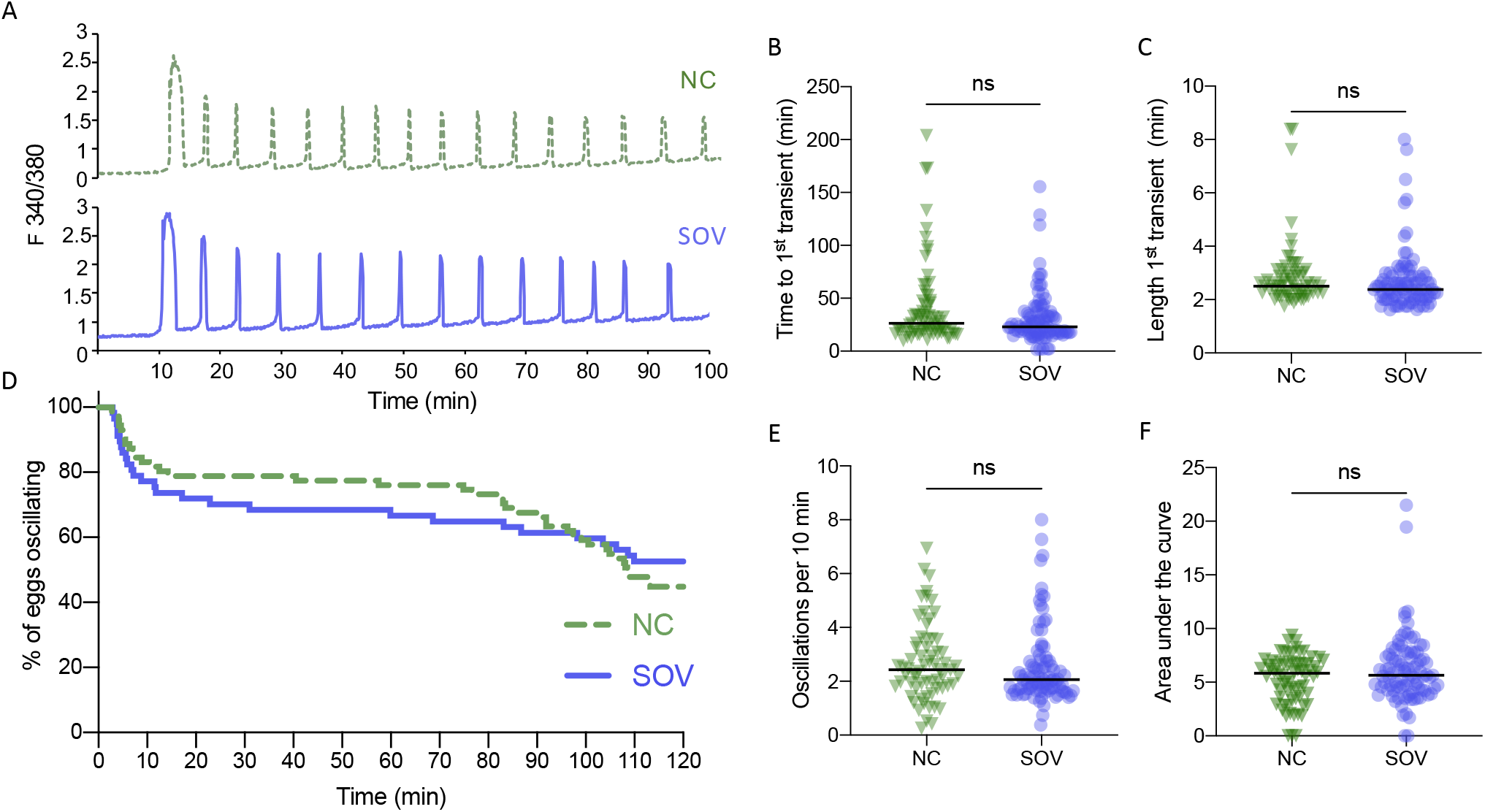
Calcium oscillatory patterns following fertilization of MII eggs from naturally cycling (NC) and superovulated (SOV) mice. (A) Representative calcium traces. (B) Time to the first transient. (C) Length of first calcium transient. (D) Percentage of eggs that continued to display calcium oscillations during 120 min. (E) Oscillation frequency averaged over the first 60 min of oscillations. (F) Area under the curve of calcium signal during the first 60 min following fertilization. Graphs in B, C, E and F show median and all individual data points. ns, no significant difference.

## Discussion

Here we tested in a highly controlled fashion whether superovulation causes abnormalities in calcium signaling following in vitro fertilization in the mouse. Despite our prediction to the contrary, the data clearly showed that in a commonly utilized inbred strain, C57BL/6J, superovulation does not alter the calcium oscillatory patterns at fertilization. Superovulation maximizes the numbers of ovulated eggs that can be obtained for study from each mouse, reducing the numbers of animals that must be euthanized for experimental purposes. Therefore, this technique is routinely performed to obtain oocytes and early embryos in laboratories and animal facilities. Our findings demonstrating normal calcium oscillatory patterns despite superovulation is reassuring regarding interpretation of experiments examining factors that influence calcium signals given that these types of experiments are almost always performed with eggs obtained by superovulation. Our data support the use of superovulation in future studies of calcium signaling at fertilization.

Despite being a widespread practice, superovulation is associated with abnormalities in several phenotypic features in oocytes and ovulated eggs. For example, an increased incidence of meiosis arrest prior to metaphase II, asynchrony between nuclear and cytoplasmic maturation, abnormalities in subcellular structure, and abnormal patterns of protein synthesis have all been observed in various mammalian species following superovulation (Moor et al., 1985; Callesen et al., 1986; Hyttel et al., 1986; Yun et al., 1989; Assey et al., 1994). Given that the hormone injections only rescue late secondary and antral follicles, these differences are clearly a consequence of inadequate preparation for oocyte maturation during later stages of follicle development. Although the oocyte is transcriptionally inactive after reaching the antral stage, translation continues to occur, creating additional maternal components to support maturation and embryo development. In addition, mRNAs and long noncoding RNAs are transferred to the oocyte from the surrounding cumulus cells, at least some of which are translated and may contribute to oocyte competence to undergo maturation (Macaulay et al., 2014, 2016). Superovulation could accelerate late follicle development enough that there is insufficient time for the transfer of these granulosa cell components and/or their translation. However, no differences were found in calcium dynamics at fertilization when SOV and NC eggs were used. These findings suggest that adequate amounts of the maternal components required for calcium homeostasis are already present prior to superovulation or, alternatively, that these components are not affected in a detrimental fashion by the superovulation protocol.

Unlike the above oocyte maturation defects, which occur prior to fertilization, some superovulation-associated abnormalities are observed following fertilization. Reduced competence to develop to or beyond the 16-cell stage, both in vivo and in vitro, abnormalities in acquisition and maintenance of methylation marks on imprinted genes and fetal growth retardation all are associated with SOV protocols in cows and/or mice (Blondin et al., 1996; Van der Auwera, 2001; Fortier et al., 2008; Market-Velker et al., 2010). Differences in embryo development likely result from a lack of accumulation of other maternal components in the final period of follicle growth to the periovulatory stage, not related to the calcium signaling toolkit. Future studies will be needed to determine the identity of such components or to develop superovulation protocols that do not disrupt preovulatory follicle development.

## Acknowledgements

We thank Briggs Hagler and Greg Scott (NIEHS) for assistance with animal handling and Safia Malki and Karina Rodriguez (NIEHS) for critical review of the manuscript. This work was supported by the Intramural Research Program of the National Institutes of Health, National Institute of Environmental Health Sciences, 1ZIAES102985.

## Author contributions

V.S., P.S., and C.J.W. designed the study. V.S. and P.S. conducted the experiments. M.S. wrote software for calcium signal analyses. V.S. analyzed the data. V.S. and C.J.W. wrote the first draft of the manuscript. V.S., P.S., M.S. and C.J.W. edited the manuscript.

